# An integrative multi-omics approach points to membrane composition as a key factor in *E. coli* persistence

**DOI:** 10.1101/2020.08.28.271171

**Authors:** Silvia J. Cañas-Duarte, Lei Sun, María Isabel Perez-Lopez, Cornelia Herrfurth, Lina M. Contreras, Ivo Feussner, Chad Leidy, Johan Paulsson, Diego M. Riaño-Pachón, Juan M. Pedraza, Silvia Restrepo

## Abstract

Many diverse bacteria can enter non- or slow-growing states where they are transiently tolerant to antibiotics. Despite its medical importance, the genetic mechanisms underlying this ‘persistence’ remain largely unknown, especially for spontaneous (type II) persistence that arise during exponential growth in rich medium. To address this challenge, here we combine genomic, transcriptomic and lipidomic analysis to identify the persistence mechanisms. We first analyzed the genome of the high-persistence mutant *Escherichia coli* DS1 (*hipQ*) to identify candidate genes for the high persistence phenotype. We then compared the gene expression profile of spontaneous persisters to normally growing cells with RNAseq and find that the activation of stress response mechanisms is likely not very important in the entrance into *hipQ*-driven spontaneous persistence. Transcriptomic results also suggest that modifications in the cell membrane play an important role, as further corroborated by lipidomic profiles showing a higher level of unsaturated fatty acids in spontaneous persisters compared to induced persisters or normally growing cells. Taken together, our results indicate that changing membrane composition is a key process in persistence, and further our understanding of spontaneous persister cells from the DS1 (hipQ) context.

## Background

Clonal populations of bacteria can stochastically generate subpopulations of slow- or non-growing cells [1–4] that are transiently tolerant to multiple antibiotics. Such ‘persister’ cells have been implicated in a wide range of chronic bacterial infections [2, 3]. Persister cells have also been associated with the recalcitrance of biofilms [5, 6]. While in biofilms, persisters are protected from the immune system by exopolymer matrices[7]. This combination of drug-tolerance and immune-evasion prevents the complete eradication of bacterial infections. Furthermore, persisters may extend infections long enough to the point where genetic resistances emerge [7, 8]. The persistence phenomena are widespread among pathogens, including *Staphylococcus aureus*, *Pseudomonas aeruginosa*, and *Mycobacterium tuberculosis* [2, 9, 10], and therefore of high public health priority.

From an ecological perspective, creating small subpopulations that survive otherwise bactericidal treatments may serve as an adaptive, bet-hedging strategy against catastrophic events [1, 11] for the population as a whole. In support of this hypothesis, some Toxin-Antitoxin (TA) loci like *hipAB* and *istR/tisB*, and stress response mechanisms have both been implicated in the generation of persister cells [12–20]. Another proposed pathway for induction of persistence is through TolC efflux pumps [21], which also display great heterogeneity across populations [22]. These findings illustrate both how external signals can influence the transition to persistence and the variety of factors that determine the state. They also have in common an important connection with the membrane. Despite these advances, the specific mechanisms of multitolerance remain largely unknown [22, 23]. The variety of insults that they can survive includes not only different types of antibiotics, but also alkaline and enzymatic lysis [24]. Furthermore, dormancy alone is not necessary or sufficient to provide persistence [25]. This indicates that there is still much to be discovered about the persistence state itself and how it results in multitolerance.

Persisters can be phenotypically classified as triggered (type I) or spontaneous (type II), depending on whether they are directly caused by stress or arise in growing populations without any known external trigger. These two persister types indeed correspond to distinctly different cell states in *E. coli* [1, 4] and seemingly involve different sets of mechanisms (Figure 1). However, either type has been hard to study because the persister state is extremely rare. Many studies have therefore relied on specific mutants that increase the persister frequency, hoping that they do not distort their wildtype counterpart in other ways. Specifically, for triggered persisters, mutant strain *hipA7* (TH1269) [1, 4, 17, 26, 27] has successfully helped to identify the involvement of TA modules and stress response mechanisms [2, 8, 17, 23, 28].

**Figure 1.**
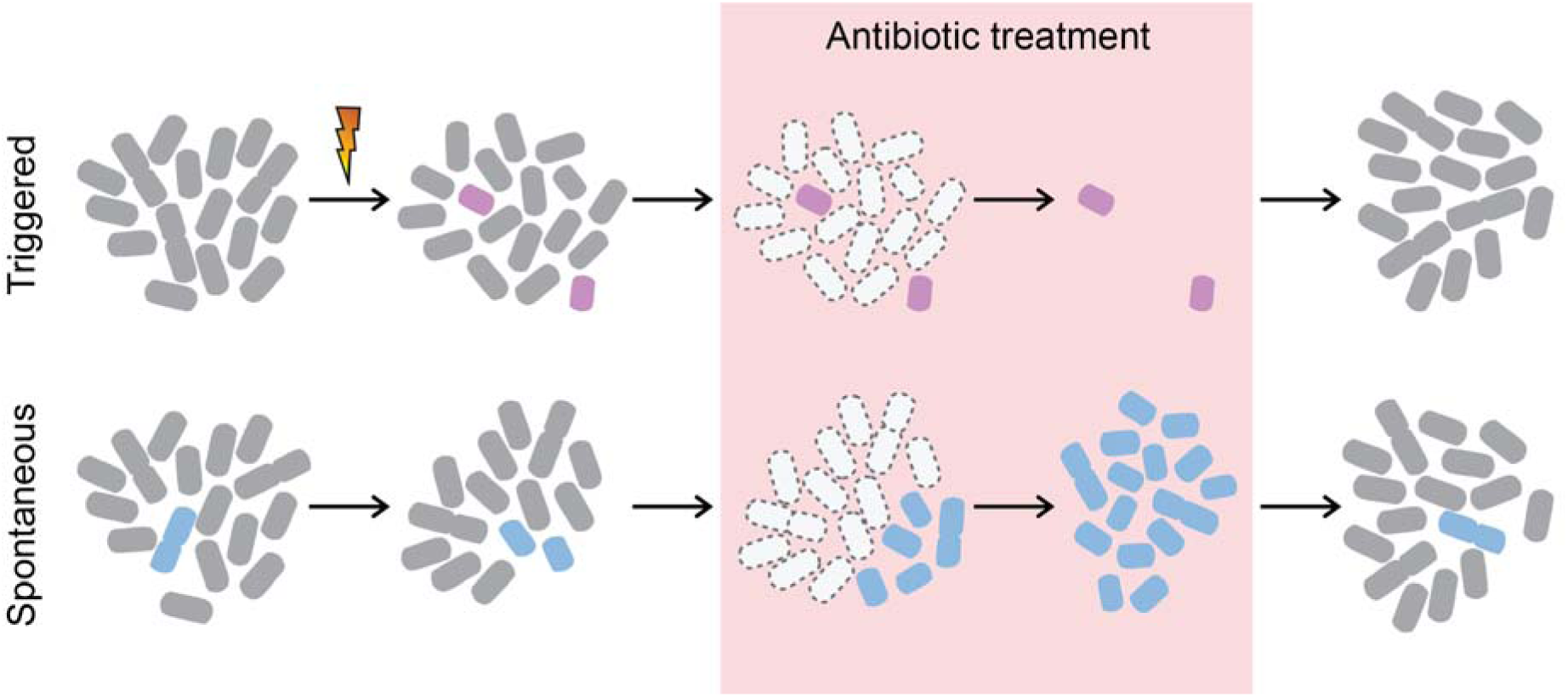
Schematic of the differences between triggered and spontaneous persisters. Triggered persisters are generated in bacterial populations upon the ocurrence of a trigger event (i.e, starvation, antibiotics, acid stress, etc) and are characteristically non-growing while in the persister state. Spontaneous persisters stochastically appear in bacterial populations, entering a slow-growing state in which they can tolerate and grow in the presence of antibiotics.

For spontaneous persisters, the high-persistence strain E. coli DS1 was isolated via mutagenesis of the parent E. *coli* strain KL16 [1, 4, 29]. By first conjugative mapping and later transducing wild-type genome fragments via P1vir into DS1 and checking for loss of high-persistence, a *hipQ* locus near the *leu* operon at the 2 min position of the DS1 chromosome was found necessary for the high-persistence phenotype of this strain [27]. However, attempts to transduce the *hipQ* phenotype to a wild-type background failed, which suggests this locus is not sufficient and that additional unidentified mutations present in DS1 genome are required [27]. Using microfluidics and microscopy, spontaneous persisters were later visualized as slow-growing cells (∼160 +/-30 min doubling-time) that arise spontaneously from an exponentially growing bacterial population [1, 24]. Entrance into this slow-growth state seems independent of antibiotics and cells showed division both before, during and after a transient antibiotic treatment. A recent study [30] presented a mutation in the *ydcI* gene (at 32 min chromosomal location) as solely responsible for the *hipQ* persister phenotype. Interestingly, these two studies contradict each other both in the gene location and number of mutations required. In addition, due to the higher difficulty in isolating spontaneous persisters [24], their overall persistence mechanism remains largely unknown.

Using our previously-published protocol capable of separately isolating triggered versus spontaneous persisters from bacterial populations [24], we performed comprehensive genomic, transcriptomic and lipidomic analyses. Genomically, we identified 59 novel missense and nonsense SNP mutations in the DS1 genome that we propose as candidates responsible for its high spontaneous persistence phenotype. Notably, we found that *ydcI* has no mutation in this strain. Furthermore, there is no DS1 strain-specific mutation near the *leu* operon where the *hipQ*-locus was proposed to be located [27]. By assessing the persister frequencies of the relevant mutants, we show that the mutations responsible for the *hipQ* phenotype of the DS1 have not yet been identified, despite previous reports. Transcriptomically, we tested whether the previously reported genetic mechanisms of persistence induction, such as stress response mechanisms, are also involved in the generation of spontaneous persister cells from the strain *E. coli* DS1 (*hipQ*) [2, 23]. Additionally, the transcriptomic analysis of spontaneous persister cells showed no evidence of SOS response activation in the spontaneous persistence physiological state, contrary to several reports [12, 13, 16, 23]. Finally, our lipidomic data suggests that modifications in the physicochemical properties of the cell membrane could be related to the formation of persister cells and constitute an important mechanism for their multitolerance to bactericidal agents. We believe these results not only broadly characterize the persister states, but also serve as a springboard for future mechanistic analyses.

## Results

### Novel SNPs identified in *E. coli* DS1 genome

To identify the genetic mechanisms related to the high spontaneous persistence phenotype of *E. coli* DS1, we first sequenced the DS1 genome, obtaining 2.5 million clean reads with a minimum length of 60 bp. The average coverage in the *de novo* genome assembly was 82,85X, with a maximum coverage of 358X, estimated using Tablet [31]. Following the de novo assembly, we obtained a single pseudo-molecule of 4567805 bp with 47 gaps and 531 N’s (Table 1). After the annotation step, 4,554 genes were predicted (Figure 2).

**Figure 2.**
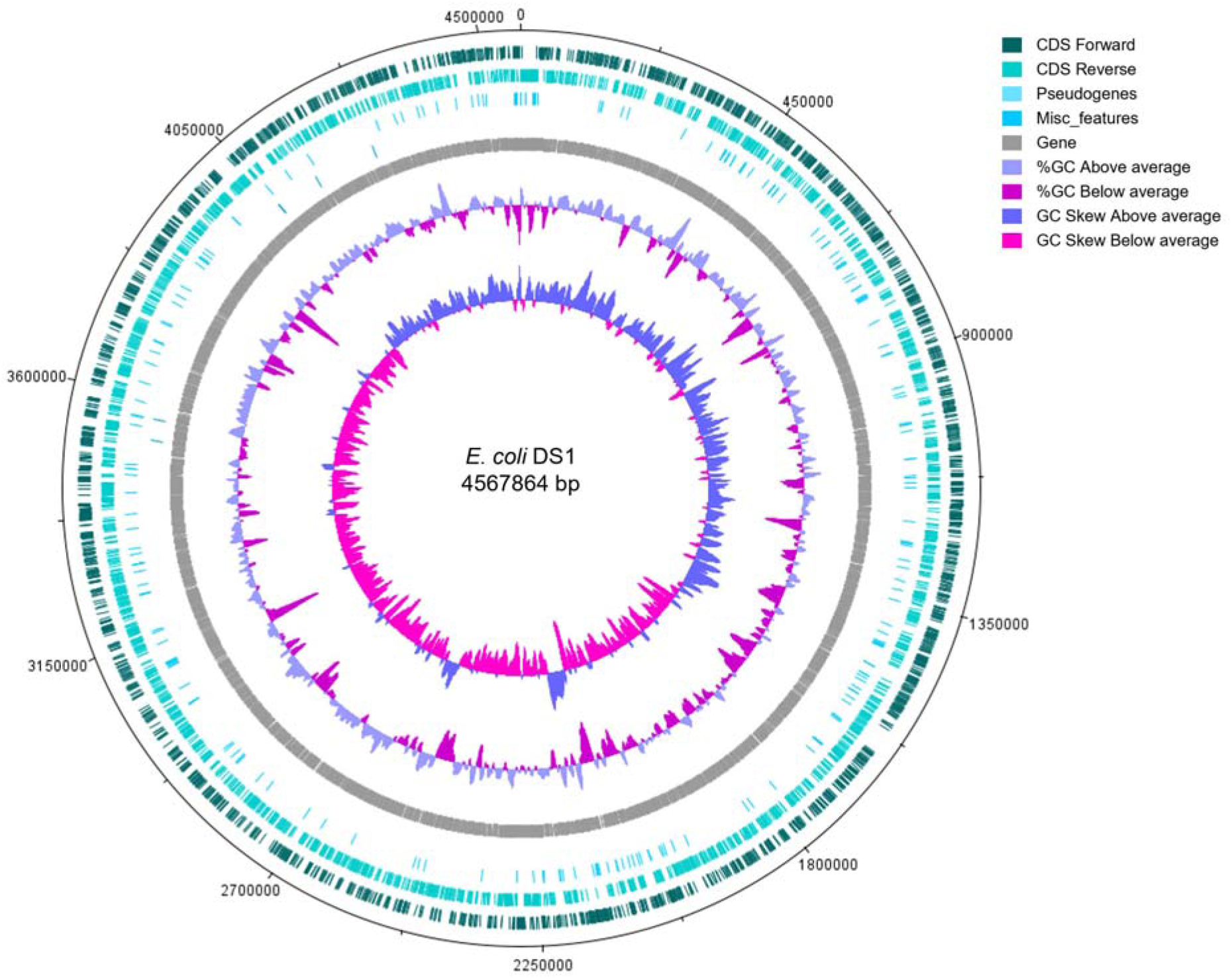
Circular representation of the novel genome assembly and annotation of *E. coli* DS1. Circular plot of the annotated DS1 genome was created using DNAplotter [36]. The tracks from the outside represent: (1) CDS Forward; (2) DDS Reverse; (3) Pseudogenes; (4) Miscellaneous features; (5) Genes; (6) %GC plot; (7) GC skew [(GC)/(G+C)]. For the %GC and GC Skew plots, a window size of 10000 bps and a step size of 200 bps were used.

**Table 1.**
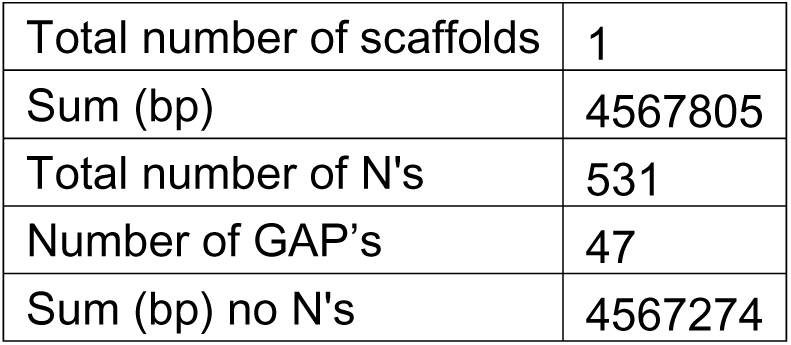
Metrics of the *E. coli* DS1 genome assembly. Assembly of the DS1 genome was performed *de novo* using CLC, obtaining initially 104 contigs. The scaffolding process was performed using first SSPACE[32] with clean PE reads from the sequencing of both the genome and transcriptome to obtain 58 scaffolds. The genome of *E. coli* MG1655 was used to refine the assembly using the PAGIT pipeline[33]. Finally, Gapfiller[34] was used to close gaps. The RAST[35] annotation server was used to predict and annotate genes in the DS1 genome.

|  |  |
| --- | --- |
| Total number of scaffolds | 1 |
| Sum (bp) | 4567805 |
| Total number of N's | 531 |
| Number of GAP's | 47 |
| Sum (bp) no N's | 4567274 |

The *E. coli* DS1 strain was evolved from the KL16 [37] parent strain (The complete genome of KLY, namely KL16 with an integrated YFP segmentation marker, is available at GenBank: CP008801.1). The comparison of KLY, MG1655 and DH10B to reference genomes revealed a total of 153, 255 and 349 SNPs, respectively (Table 2). As none of the reference strains show a high persistence phenotype, and the DH10B strain presents persistence frequencies below the wild type levels (data not shown), we only considered the 112 novel SNPs that were detected against all three reference genomes for further analyses (Additional file 1). The genomic analysis performed also showed the presence of the F plasmid integrated into the chromosome [27, 29].

**Table 2.** Analysis of Single Nucleotide Polymorphisms (SNPs) detected in the genome of *E coli* DS1. The genomes of the KLY (KL16 with YFP) parental strain, the wild type strain MG1655 and the DH10B strain were used as a reference; we detected a total of 153, 255 and 349 SNPs, respectively.

| Type | Strain | Count | Percent/ (Total SNPs) |
| --- | --- | --- | --- |
| MISSENSE | MG1655 | 132 | 43.40% |
|  | KLY | 58 | 37.90% |
|  | DH10B | 122 | 73.05% |
| NONSENSE | MG1655 | 3 | 1.57% |
|  | KLY | 1 | 0.65% |
|  | DH10B | 3 | 1.80% |
| SILENT | MG1655 | 109 | 52.60% |
|  | KLY | 89 | 58.17% |
|  | DH10B | 224 | 64.20% |

As its parent strain KL16, E. coli DS1 encodes the *relA1* and *spoT1* mutations, along with several other polymorphisms when compared to the MG1655 strain. It has been previously reported that strains with *relA1* and *spoT1* mutations have deficient stringent response [38]. The wild-type persistence levels of the KL16 strain and the fact that the *hipQ* phenotype of DS1 is most relevant during exponential phase and conditions free of external stressors, strongly indicate that these mutations are not directly involved in the *hipQ* phenotype.

Some of the identified polymorphisms appear on genes whose functions were previously reported to be related to persistence, such as stress response mechanisms, DNA replication and catabolic processes of amino acids and carbon sources [8, 17]. We found a novel mutation in the *hipA* locus (A242V) different from the previously characterized hipA7 mutations known to confer a high persistence phenotype [17]. However, this mutation, although not generally considered a conservative change, does not appear to generate any significant conformational change in the protein (Additional file 2), although it is located near the active site of this kinase. We also identified various mutations in genes involved in the metabolism and transport of lipids such *fadD, ytfN (tamA)* and *ytfM* (*tamB*), cell wall homeostasis such murB, *yceG* (*mltG*) and *amiB*, and transmembrane transport such as *ompF and ompN*. As discussed in the next section, these results suggest the existence of modifications in cell envelope physicochemical properties in persister cells, as several components related to the cell envelope were found to be overrepresented amongst the set of genes in which novel mutations were identified for DS1 (Table 3). As noted above, our genomic analysis found no mutations in the gene *ydcI*, recently reported as the gene responsible for the high spontaneous phenotype persistence [30].

**Table 3.** Overrepresentation analysis of the Cellular components represented in the set of genes with novel SNPs in DS1. To analyze which cellular components were being significantly overrepresented amongst the set of mutated genes in DS1, PANTHER [39, 40] was used to map the genes to UniProt accession numbers. DAVID [41, 42] then used to test for overrepresentation of cellular components, using as reference the complete genome of E. coli K12.

| Go term<br>(Cell component) | Count | p-value | Fold<br>enrichment | Fisher Exact<br>test |
| --- | --- | --- | --- | --- |
| Cell envelope | 18 | 1.1E-03 | 2.3 | 4.3E-04 |
| Outer membrane-bounded periplasmic<br>space | 11 | 1.8E-02 | 2.3 | 7.5E-03 |
| Intracellular organelle<br>lumen | 3 | 2.5E-02 | 11.7 | 1.8E-03 |
| Organelle lumen | 3 | 2.5E-02 | 11.7 | 1.8E-03 |
| Membrane-enclosed<br>lumen | 3 | 2.5E-02 | 11.7 | 1.8E-03 |
| Membrane | 45 | 3.0E-02 | 1.2 | 2.3E-02 |
| Periplasmic space | 12 | 3.2E-02 | 2 | 1.5E-02 |
| Intracellular membrane-bounded organelle | 4 | 4.2E-02 | 5 | 7.5E-03 |
| Membrane-bounded<br>organelle | 4 | 4.5E-02 | 4.9 | 8.3E-03 |
| 2-iminoacetate synthase complex | 2 | 4.6E-02 | 42.8 | 5.4E-04 |
| TAM protein secretion complex | 2 | 4.6E-02 | 42.8 | 5.4E-04 |
| Pore complex | 3 | 7.0E-02 | 6.8 | 9.1E-03 |
| Cell outer membrane | 7 | 8.2E-02 | 2.3 | 3.3E-02 |
| Outer membrane | 7 | 9.9E-02 | 2.2 | 4.2E-02 |

To further corroborate this, we designed specific primers for the genes *ydcI* and *ybaI* and performed Sanger sequencing. Again, no mutations were detected in these genes. Furthermore, we tested the persister frequency to ampicillin of a Δ*ydcI* strain generated by P1 transduction from the KEIO collection [43] into MG1655. According to the previous report [30], the MG1655 Δ*ydcI* strain should show a similarly high persister frequency as that of the DS1. Our results, however, show this is not the case, with this strain having persister levels close to that of the MG1655 wild-type, even slightly lower (Figure 3). In addition, we searched for mutations located within ∼100 kb from the *leu* operon, the proposed site of the *hipQ* locus. To our surprise, DS1 has no strain-specific point mutations in the said region. Instead, it contains two SNPs inherited from its KL16 parent strain: *creC* R77P and *aceE* A20T. To assess whether these two mutations contribute to high-persistence, we restored each of these mutations independently with P1 transduction using the respective wild-type gene from the MG1655 strain and moved it into the DS1 strain and then tested the persister frequency to ampicillin. For both the DS1 *creC* and DS1 *aceE* restored strains, we found no significant changes in their persister frequencies when compared to the original DS1 strain, indicating that this mutations inherited from KL16 have no effect on the hipQ phenotype (Figure 3). So far, our results suggest that all previous findings on the *hipQ* genetics are likely incorrect. Therefore, we conclude that the genetic mechanism of *hipQ* is not yet determined and remains an open question.

**Figure 3.**
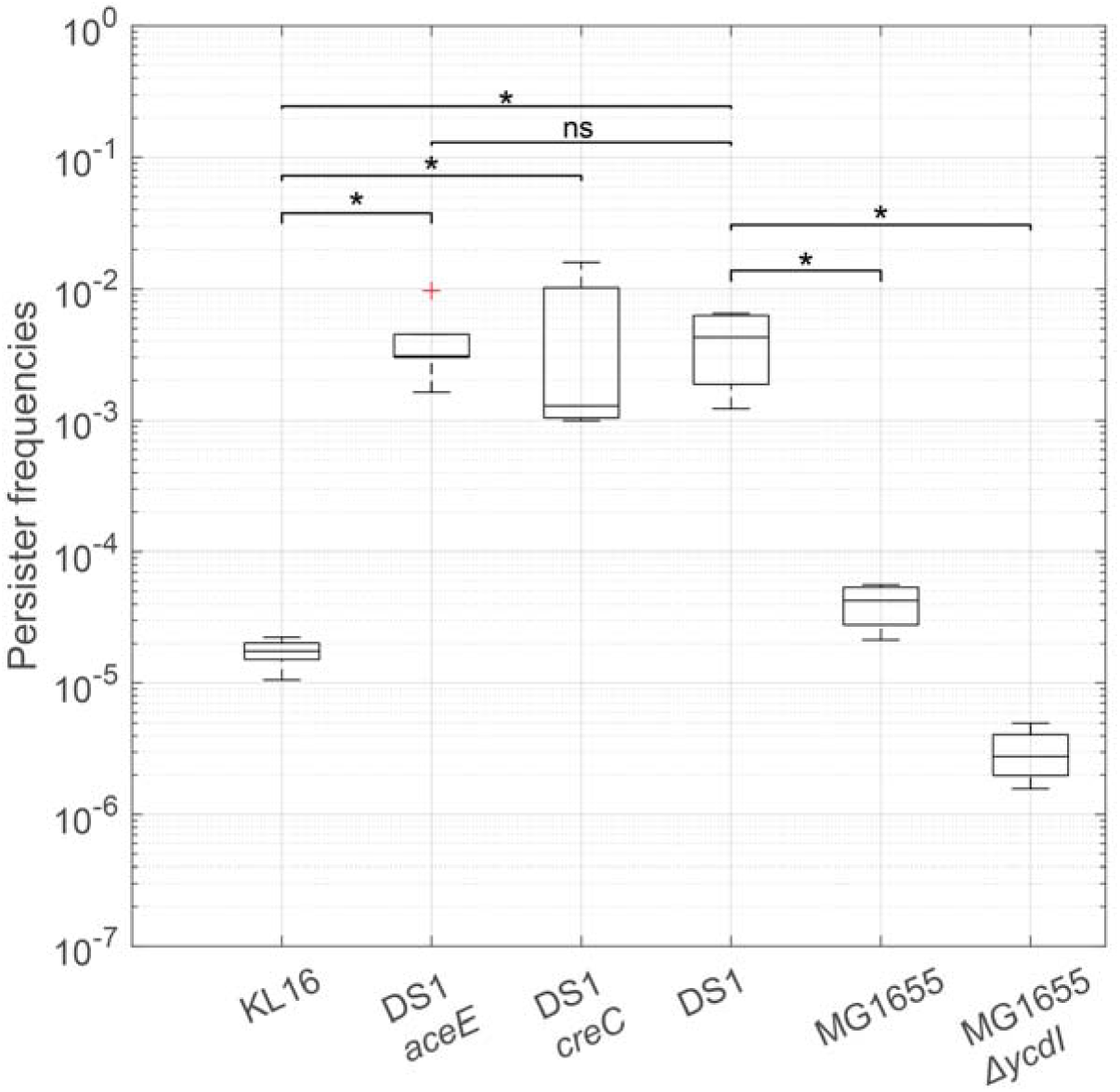
Comparison of persister frequencies of previously reported *hipQ* related mutations in the wild-type strains (KL16 and MG1655) and the *hipQ* (DS1) strain. Persister frequencies were determined by ampicillin treatment for 3 hours, as described previously [1]. Ampicillin (100 ug/mL) was added to the growing culture when it reached an OD600 = 0.4. Time 0 (before addition of antibiotic) and t=3 hours were serially diluted and plated in LB agar to determine CFUs. Values represent the average of 3 biological replicates. Two-sample t-test were performed to determine if the differences between the persister frequencies the indicated pairs of strains are significant (p-value < 0.05).

### Differentially expressed genes in spontaneous persisters were distinct from previously reported transcriptomes of persister cells

Due to the large number of SNPs detected, pinpointing the mechanism with genome alone is challenging, especially since multiple mutations are likely involved [27]. Therefore, we carried out transcriptomic analysis. Previously, persisters have been routinely isolated by antibiotic treatments that lasts a few hours [1, 8, 10, 16]. Alternatively, triggered persister studied have used flow cytometry to enrich persisters by using fluorescent reporters of ribosomal genes like *rrnbP1* [44]. Depending on the choice of antibiotics, cell physiology could be differentially affected [16, 45]. Since spontaneous persisters emerge independently of antibiotic treatment, and to avoid antibiotic-induced biases, we used a previously published lysis protocol that isolates persisters in under 30 min (Additional file 3). From the isolated DS1 persisters, we performed transcriptomic analysis and compared to the transcriptome of exponentially growing, non-persister cells of DS1.

Utilizing the paired end and strand-specific reads obtained during the RNASeq analysis, we assembled a complete gene catalog of expressed genes of *E. coli* DS1 cells during spontaneous persistence and exponential growth using 27,525,259 and 40,130,287 paired end reads, respectively, with an average length of 78 bps. For spontaneous persister cells, we assembled and annotated 5,768 transcripts; 5,748 of these transcripts displayed BLAST hits, whereas the number of transcripts was 4,393 for the exponentially growing cells.

A total of 301 statistically significant differentially expressed genes (DEG) were detected. Of these, 217 were found to be down-regulated and the remaining 84 up-regulated. In the list of up-regulated genes, we noticed the overall absence of genes related to stress response mechanisms that were previously reported to be involved in the generation of persister cells [8, 22, 28]. Notably, genes participating in the SOS mechanism did not display differential regulation during spontaneous persistence. Several genes related to other stress-response mechanisms were found in the list of down-regulated genes (Additional file 4).

Seven genes involved in lipid biosynthesis and modifications were found to be differentially regulated during spontaneous persistence, such FadB, along with genes involved in the regulation of other cell envelope components such peptidoglycan and LPS. This is consistent with the identification of 9 novel mutations in genes related to lipid pathways in the DS1 genome. 16 genes from TA modules were found to be differentially expressed during this type of persistence. But their behavior varied across TA modules: some were down-regulated, such as, *hipB, dinJ* and *relB,* whereas others, such as *pspB*, were up-regulated. In particular, we found that the *tisB* toxin was down-regulated.

### Lipid metabolism is associated with spontaneous persistence

We analyzed genes that were differentially expressed to detect whether some biological functions were significantly over- or underrepresented in this cluster. We found that genes related to regulation of translation, regulation of protein metabolic processes, several macromolecules biosynthetic processes, response to stimuli and response to stress were significantly overrepresented in the differentially expressed gene cluster, highlighting the importance of the regulation of these functions in spontaneous persister cells.

We separately analyzed the clusters of up-regulated and down-regulated genes to identify the biological functions and processes related to spontaneous persistence. In the down-regulated gene cluster during spontaneous persistence, we found that genes related to cell division, responses to stimulus and stressful conditions, synthesis of macromolecules, translation and overall homeostasis related processes were significantly overrepresented (Figure 4B).

**Figure 4.**
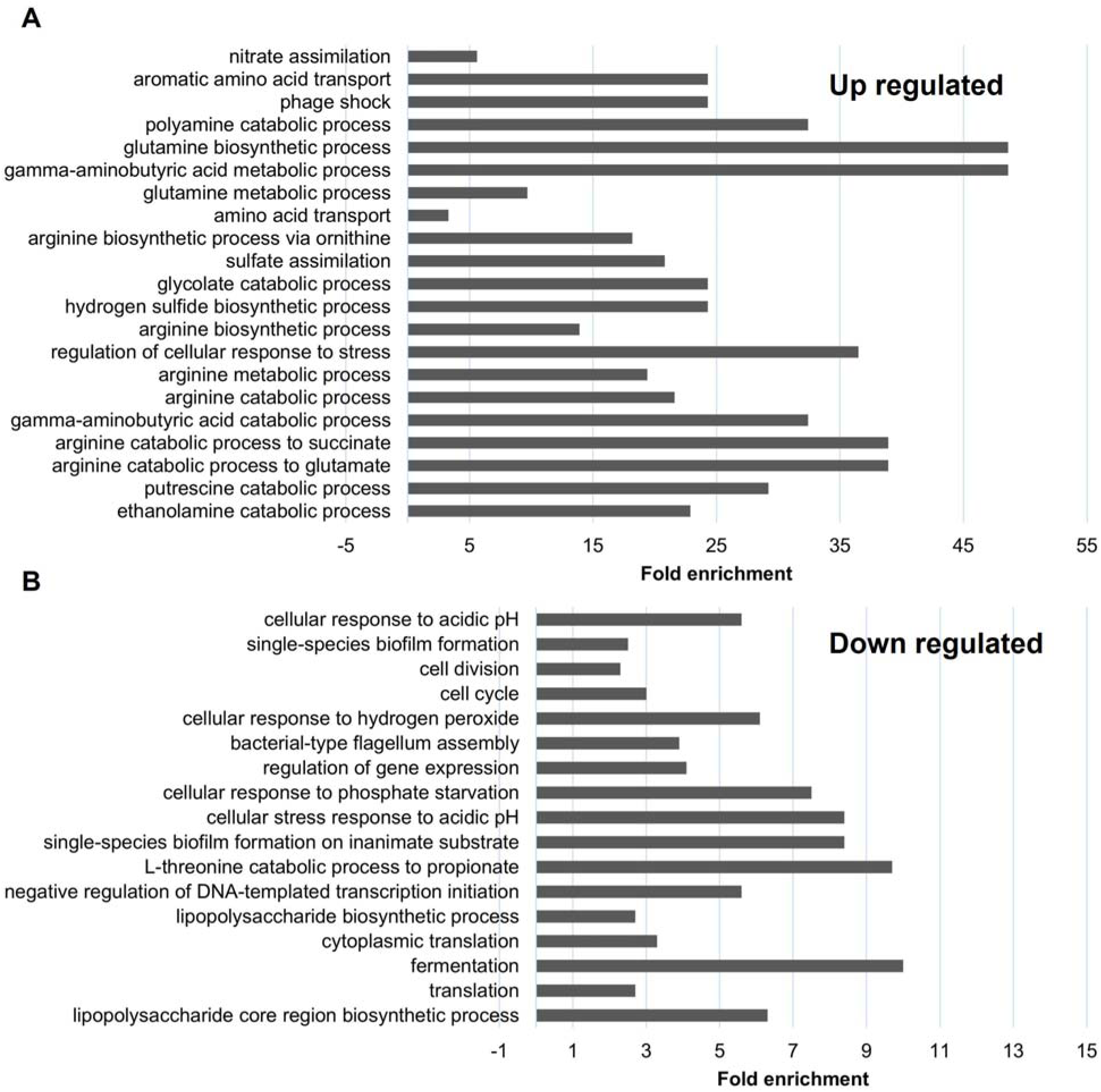
Biological processes significantly overrepresented during spontaneous persistence. To analyze which biological processes being significantly regulated during spontaneous persistence, Blast2GO[46] was used to map all the genes from the subsets of Upregulated and Downregulated DEGs. DAVID [41, 42] was then used to test for overrepresentation of biological functions in the subsets of A) up-regulated and B) down-regulated. As noted, several processes involving both DNA and RNA binding and metabolism of proteins are overrepresented in the set of down-regulated genes in spontaneous persister cells, whereas the metabolism of cell envelope components is overall overrepresented in both the up-regulated and the down-regulated clusters, indicating a strong regulation occurring during spontaneous persistence in this function. Only biological functions found to be overrepresented with a Bonferroni score < 0.005 are shown and an FDR < 0.0005.

When analyzing the cluster of up-regulated genes, we found that functions related to the intake of carbon sources, polyamine metabolism and to general catabolic processes were overrepresented. Biological functions related to lipid metabolic processes were significantly overrepresented (i.e, ethanolamine catabolic process), suggesting that modifications in the cell membrane composition and physicochemical properties are important during spontaneous persistence (Figure 4A).

### RT-qPCR validates all tested differentially expressed genes predicted by our RNASeq analysis

We selected a subset of 12 genes (Additional file 5) out of 301 total genes found to be differentially expressed with the RNASeq analysis for validation: 4 and 8 from the up- and down-regulated cluster, respectively. Besides the differential expression, the selection criteria for this subset of genes included biological functions and their relationship with mechanisms previously related to the persistence phenomenon [8, 22, 28].

To select appropriate reference genes for the RT-qPCR experiments, the expression profile of previously reported housekeeping genes in the RNASeq dataset was analyzed. The expression of the genes *opgH* and *dxs* appeared to be suitable as RT-qPCR reference genes, whereas other commonly used housekeeping genes in *E. coli*, such as *recA* and *mdh*, showed variations in their abundances when comparing the *E. coli* DS1 strain during its spontaneous physiological state to its exponential growth.

All tested genes displayed similar expression profiles in the qPCR analysis, as reported by our RNASeq analysis (Additional file 6, including the housekeeping genes *opgH* and *dxs* (data not shown).

### Identification of genes with non-synonymous mutations related to gene expression changes in E. coli DS1 spontaneous persisters potentially responsible for its high persistence phenotype

Firstly, we identified 19 SNPs directly related to the differential expression of genes in spontaneous persistence (Additional file 7). Amongst these differentially expressed genes a non-synonymous mutation in *narZ*, which encodes for the nitrate reductase Z subunit α, is related with the up-regulation of the NarZWY operon. Significantly, we found that a non-synonymous mutation in the transcriptional terminator Rho correlates with the differential expression of at least 16 genes from the putative Rho-dependent termination regulon [47]. The remaining 17 mutations identified exhibit correlations between regulators like SoxR and the mutated genes.

We then decided to directly assess the gene expression profiles of the genes found to be mutated in the DS1 genome. For this, we started by focusing on the cluster of 57 unique genes in which at least one non-synonymous SNP was detected. From this cluster, we identified 18 genes with a log2 Fold change >+/-1 in gene expression from the differential expression analysis of spontaneous persisters compared to non-persisters (Table 4). We then analyzed each of the mutated genes in terms of biological function, and assessed their possible relevance to persistence by combining their gene expression changes in persister cells with several reported factors involved in persistence such as stress response mechanisms, DNA replication and cell division and catabolic processes of amino acids and carbon sources [8, 17, 28]. With this functional analysis, we selected 10 additional genes carrying non-synonymous mutations that we consider might be potentially involved in *hipQ*-high persistence but that do not exhibit large expression changes (Table 4), whose functions we discuss in the Discussion section below.

**Table 4.**
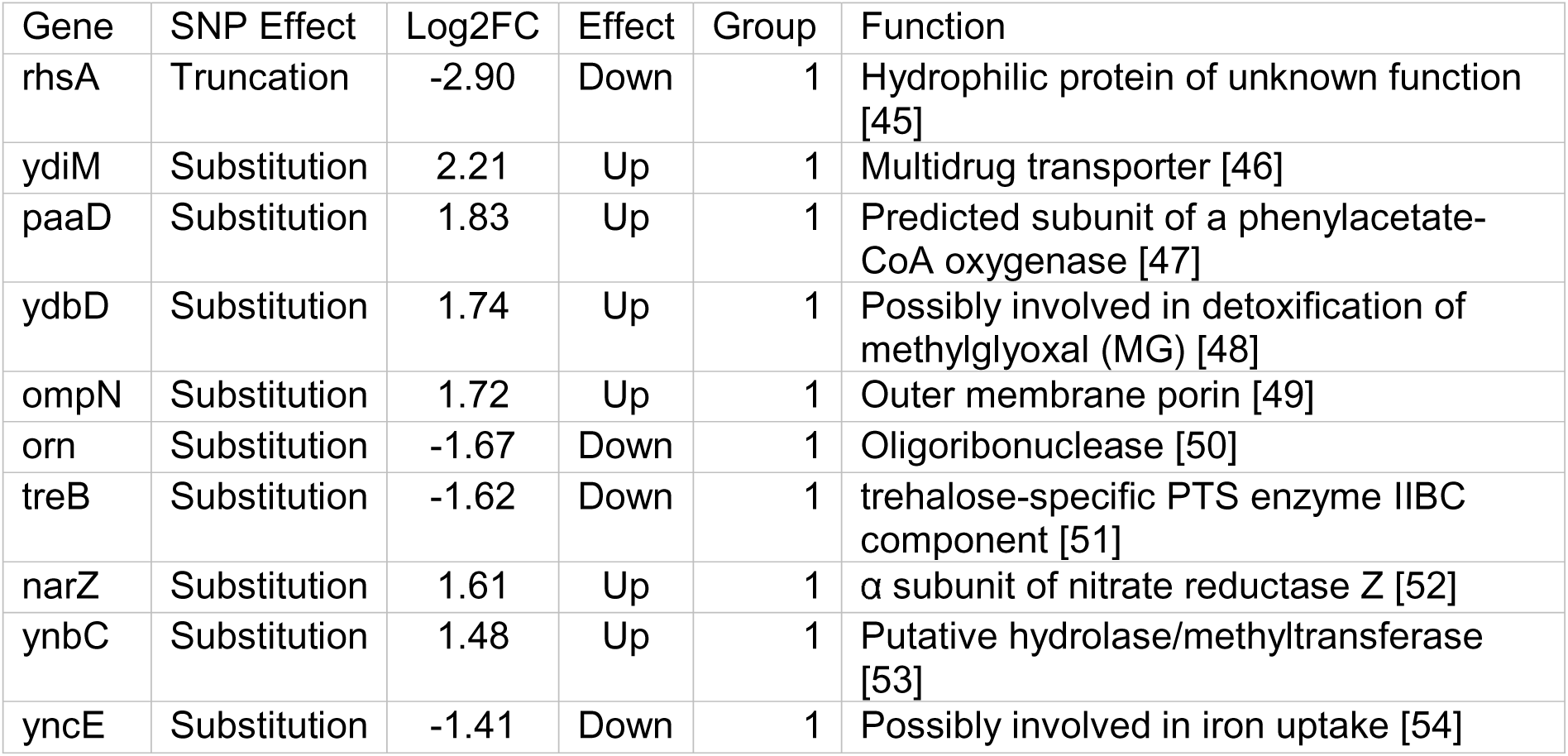
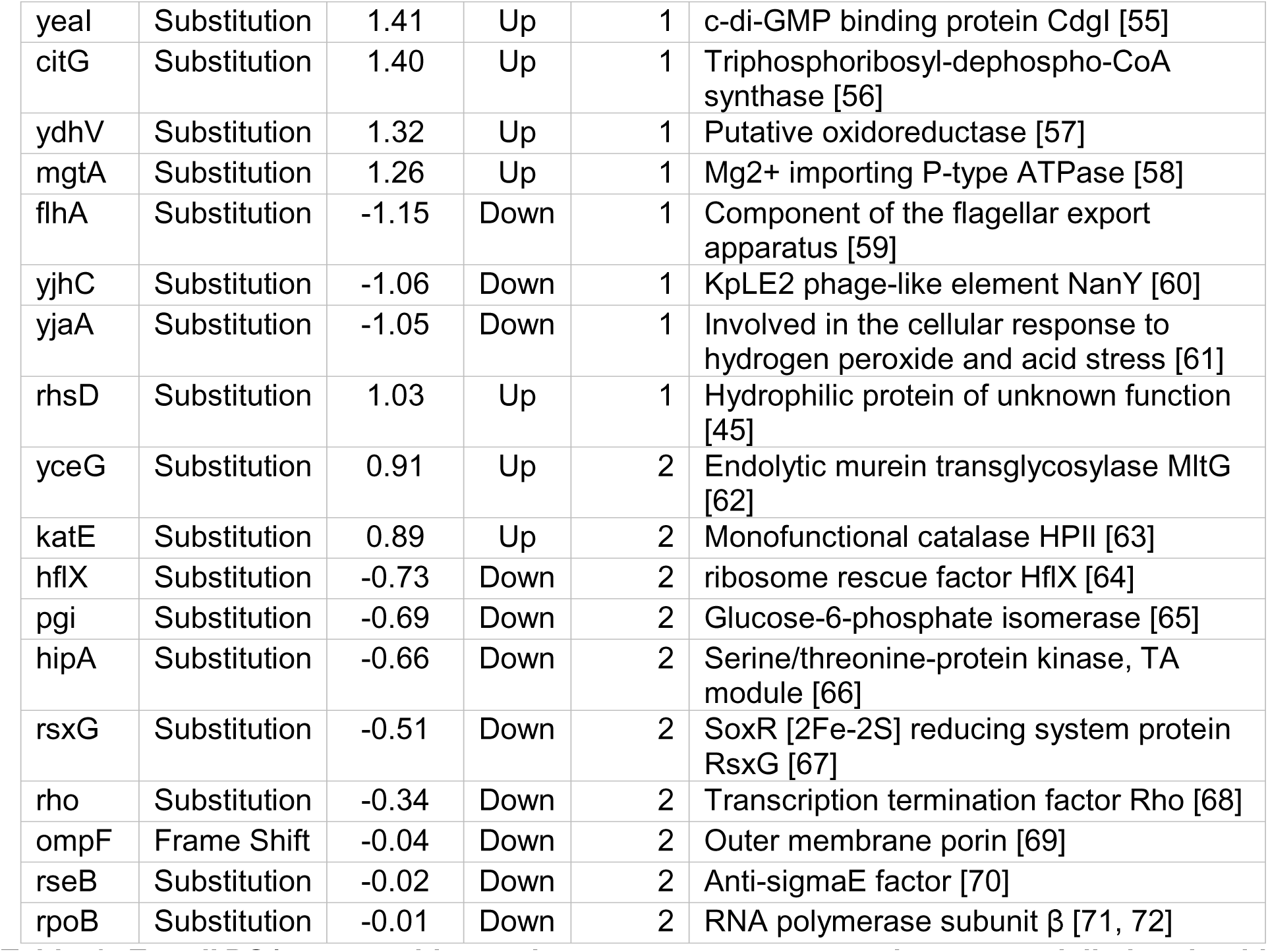
*E. coli* DS1 genes with novel non-synonymous mutations potentially involved in *hipQ*-driven persistence. 18 genes were selected with changes in expression > 2-fold in spontaneous persisters compared to the exponentially growing cells from which they emerged (Group 1); 10 additional genes were selected based on their biological function having been linked to persistence but with smaller or undetected changes in expression (Group 2).

| Gene | SNP Effect | Log2FC | Effect | Group | Function |
| --- | --- | --- | --- | --- | --- |
| *rhsA* | Truncation | -2.90 | Down | 1 | Hydrophilic protein of unknown function [45] |
| *ydiM* | Substitution | 2.21 | Up | 1 | Multidrug transporter [46] |
| *paaD* | Substitution | 1.83 | Up | 1 | Predicted subunit of a phenylacetate-CoA oxygenase [47] |
| *ydbD* | Substitution | 1.74 | Up | 1 | Possibly involved in detoxification of methylglyoxal (MG) [48] |
| *ompN* | Substitution | 1.72 | Up | 1 | Outer membrane porin [49] |
| *orn* | Substitution | -1.67 | Down | 1 | Oligoribonuclease [50] |
| *treB* | Substitution | -1.62 | Down | 1 | trehalose-specific PTS enzyme IIBC component [51] |
| *narZ* | Substitution | 1.61 | Up | 1 | $\alpha$ subunit of nitrate reductase Z [52] |
| *ynbC* | Substitution | 1.48 | Up | 1 | Putative hydrolase/methyltransferase [53] |
| *yncE* | Substitution | -1.41 | Down | 1 | Possibly involved in iron uptake [54] |
| yeal | Substitution | 1.41 | Up | 1 | c-di-GMP binding protein Cdgl [55] |
| citG | Substitution | 1.40 | Up | 1 | Triphosphoribosyl-dephospho-CoA synthase [56] |
| ydhV | Substitution | 1.32 | Up | 1 | Putative oxidoreductase [57] |
| mgtA | Substitution | 1.26 | Up | 1 | Mg<sup>2+</sup> importing P-type ATPase [58] |
| flhA | Substitution | -1.15 | Down | 1 | Component of the flagellar export apparatus [59] |
| yjhC | Substitution | -1.06 | Down | 1 | KpLE2 phage-like element NanY [60] |
| yjaA | Substitution | -1.05 | Down | 1 | Involved in the cellular response to hydrogen peroxide and acid stress [61] |
| rhsD | Substitution | 1.03 | Up | 1 | Hydrophilic protein of unknown function [45] |
| yceG | Substitution | 0.91 | Up | 2 | Endolytic murein transglycosylase MltG [62] |
| katE | Substitution | 0.89 | Up | 2 | Monofunctional catalase HPII [63] |
| hflX | Substitution | -0.73 | Down | 2 | ribosome rescue factor HflX [64] |
| pgi | Substitution | -0.69 | Down | 2 | Glucose-6-phosphate isomerase [65] |
| hipA | Substitution | -0.66 | Down | 2 | Serine/threonine-protein kinase, TA module [66] |
| rsxG | Substitution | -0.51 | Down | 2 | SoxR [2Fe-2S] reducing system protein RsxG [67] |
| rho | Substitution | -0.34 | Down | 2 | Transcription termination factor Rho [68] |
| ompF | Frame Shift | -0.04 | Down | 2 | Outer membrane porin [69] |
| rseB | Substitution | -0.02 | Down | 2 | Anti-sigmaE factor [70] |
| rpoB | Substitution | -0.01 | Down | 2 | RNA polymerase subunit $\beta$ [71, 72] |

### Analysis of fatty acid profiles validates the occurrence of significant membrane modifications in persister cells

The results of our RNA-Seq analysis of triggered and spontaneous persisters show that genes involved in fatty acid biosynthesis, normally involved in the modulation of fatty acid chemical structure, become differentially expressed, indicating a possible role of lipid composition of the membrane during persistence. Furthermore, our previous observations showed that each type of persister tolerates different concentrations of a lytic cocktail designed to induce high osmotic stress and cell wall degradation [24]. This led us to study the lipid composition of the different types of persisters.

Overall, the fatty acid composition in normally growing cells from different *E. coli* strains is highl conserved. We also only observe minimal variations among the different physiological states for most cases (Figure 5). However, significant differences were found when comparing both triggered and spontaneous persister cells with all other samples from non-persister cells during both the exponential growth and stationary phases (Figure 5 and Additional file 8). In triggered persisters, the proportion of saturated lipids changed significantly, increasing to more than 68% in these persisters cells from less than 50% in non-persister stationary phase cells. This change is expected to significantly increase the membrane rigidity of triggered persisters, which could be related with its non-growing phenotype. In contrast to the non-growing triggered persisters, spontaneous or Type-II persisters have been shown to be dividing, albeit with a prolonged division time of ∼160 +/-30 min [1, 24]. Interestingly, spontaneous persisters increase the proportion of unsaturated fatty acids in their membrane, from an average of 41% in non-persisters to more than 58% in spontaneous persister cells, suggesting an overall increase in membrane fluidity of the cell membrane for spontaneous persisters. No changes in the average fatty acid chain length were observed for neither triggered nor spontaneous persister cells compared to normal cells (Figure 6)

**Figure 5.**
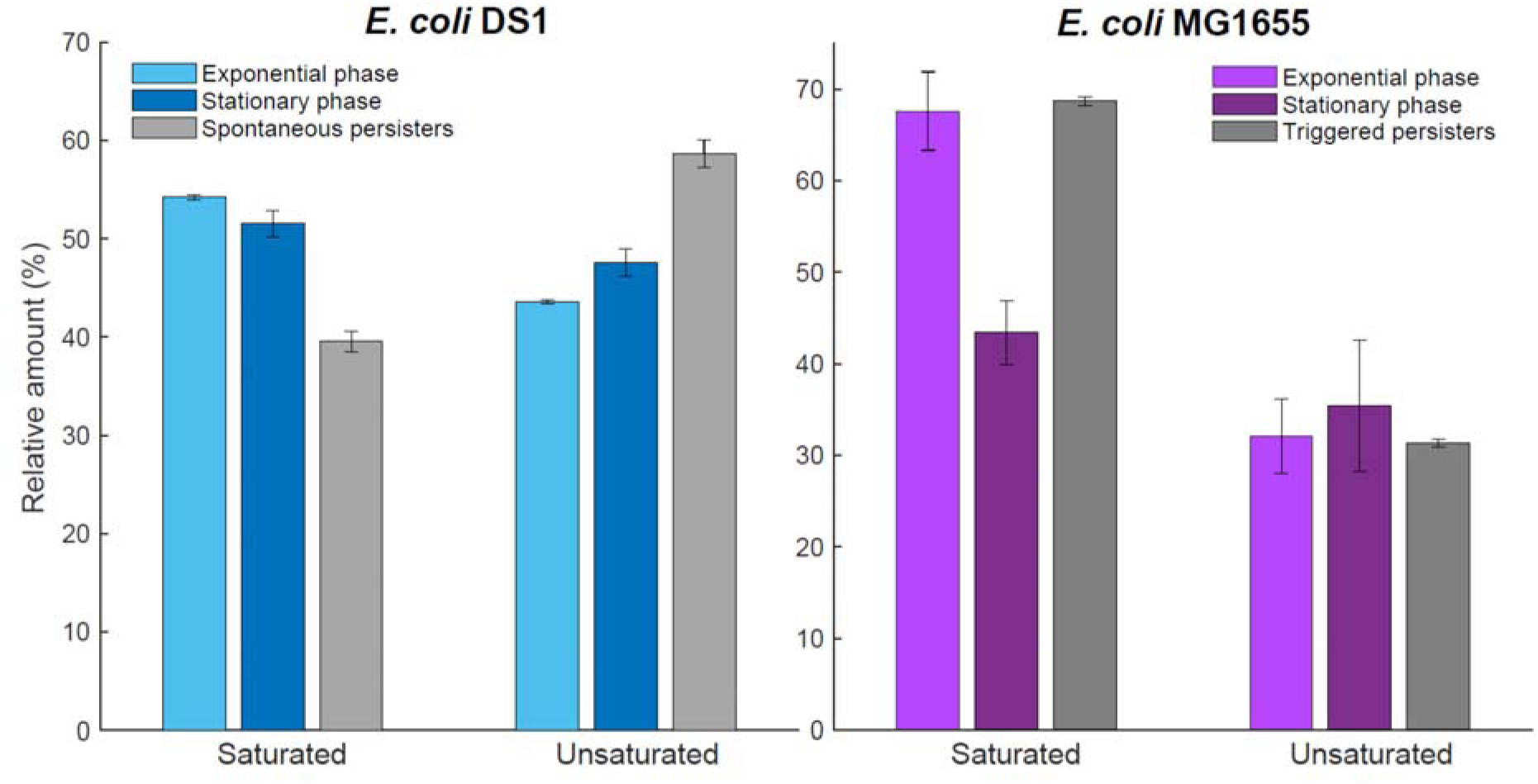
Fatty acid chain profiles evidence the occurrence of membrane modifications in triggered and spontaneous persisters. The lipid composition of persister cells is markedly different from that of the cells in the physiological state in which they are generated. (right) Triggered persisters, generated in stationary phase, significantly increase their membrane rigidity by increasing the saturated lipids at the expense of reducing the amount of unsaturated lipids, whereas the membranes of (left) spontaneous persisters, generated during exponential growth, are highly enriched in unsaturated lipids, consistent with an overall increase in the fluidity (decrease in lipid packing) of the cell membrane in persister cells.

**Figure 6.**
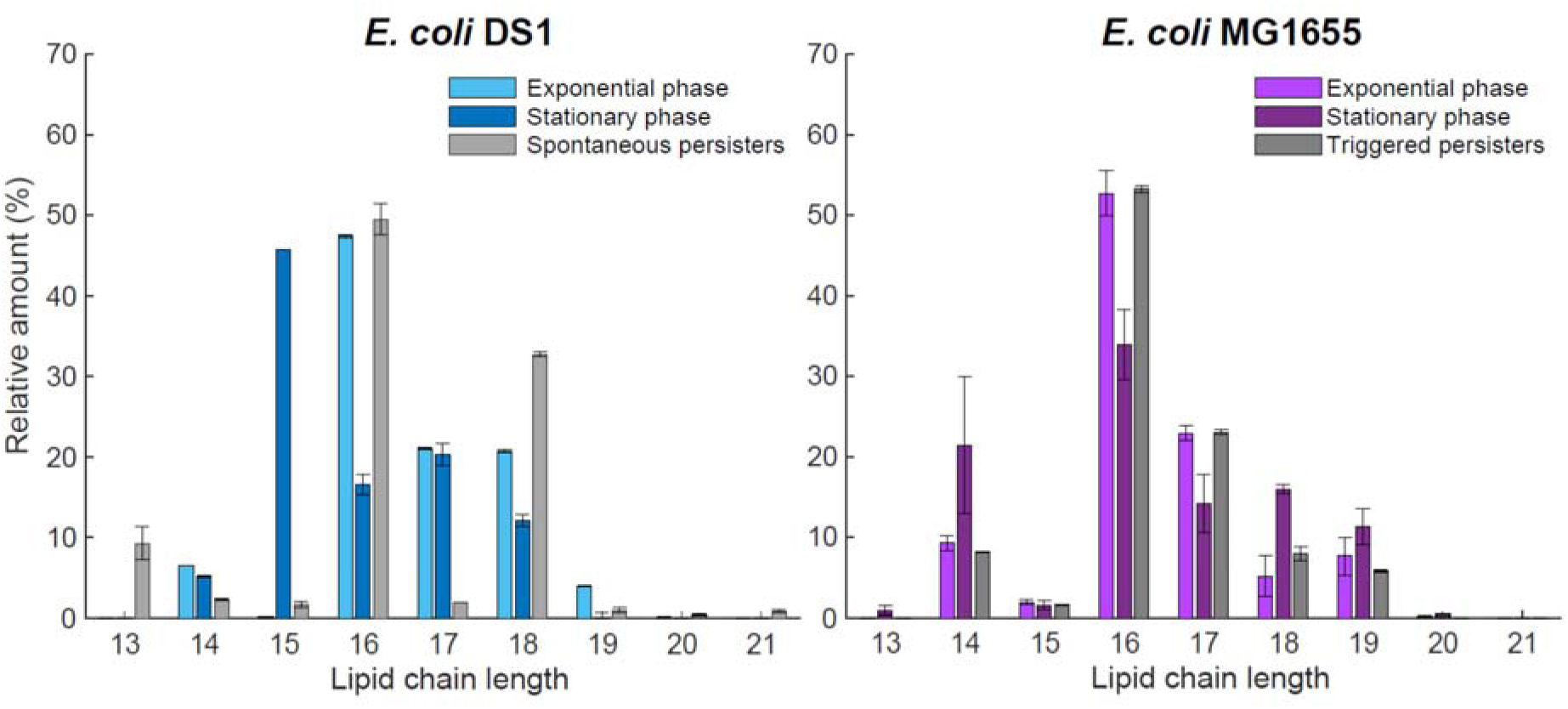
The fatty acid chain-length profiles of persisters are distinct from normally growing cells. The chain length of the lipids composing cell membranes of triggered and spontaneous persisters exhibit noticeable changes in composition. Triggered persisters (right) are enriched in both the 16 and 17 carbon chains, consistent with an increase in the saturated version of this species, compared with cells in stationary growth. Spontaneous persisters (left) are enriched in 18 carbon chains compared with cells in exponential growth, which is consistent with an increase in the unsaturated 18:1Δ9 species. The average chain length in both triggered and spontaneous persisters does not exhibit a significant change compared to normally growing cells, which is highly conserved amongst different physiological states and different *E. coli* strains.

## Discussion

Despite a great and long-standing interest in bacterial persistence, much is still unknown about the underlying biochemical and molecular mechanisms by which they are generated as persister cells are too infrequent and their transient nature difficult to study. Recent consensus has categorized persisters largely into two types: triggered versus spontaneous, previously known as type 1 versus type 2 [4] . Persistence could also arise from other mechanisms such as balanced division and deaths under antibiotic treatment [48]. Furthermore, host tissues could cause phenotypic variations of bacterial pathogens and contribute to persistence[49] .

For induced persisters, a great deal has been learned from the high persistence strain *hipA7*, mutated in the gene for the HipA toxin. For spontaneous persisters, *hipQ* has played a similar role in phenotypic analyses, as the only published mutant to our knowledge that substantially increases the spontaneous persistence rate. However, as opposed to *hipA*, the *hipQ* designation does not refer to any known gene. A major early attempt using conjugative mapping [27] failed to identify specific genes, and though a recent paper suggested a gene [30], that polymorphism does not appear to exist in this strain (the DS1 *hipQ* strain). Starting with improved isolation methods for separating triggered and spontaneous persister cells [24], we here present a multi-omic approach to the classic *hipQ* persister strain, and for the first time identify not only the key polymorphisms, but also expression profiles and lipid modifications.

Our analysis of the complete *E. coli* DS1 genome identified 121 polymorphisms unique to this strain, including a cluster of 57 genes with non-synonymous mutations potentially involved in the generation of persister cells. One notable mutation found in the DS1 genome is a novel polymorphism in the coding sequence of the HipA toxin gene, a single point mutation changing Alanine for Valine (A242V). This mutation was distinct from the hipA7 mutation (G22S+D291A) responsible for the hip phenotype of hipA7 strain [17, 26, 50]. However, we do not expect the A242V mutation to cause significant conformational changes to the HipA protein, since alanine and valine are structurally and functionally very similar. Several additional mutations occurred in genes previously implicated in persistence in *E. coli* cells, such as for genes related to stress response mechanisms, biofilm formation, quorum sensing and catabolic processes [6, 8, 10, 13, 17, 28, 51, 52]. Notably, the analysis of the DS1 genome supported the hypothesis that modifications to the lipids composing the cell membrane might influence or cause multitolerance.

We further sequenced the complete transcriptome of spontaneous persisters using exponentially growing cells as a control. A total of 180 genes were differentially expressed during spontaneous persistence: 64% were down-regulated, and 36% were up-regulated. Many genes related to carbohydrate metabolism were up-regulated, whereas genes encoding for cell division, protein synthesis and cell homeostasis processes were down-regulated. This regulation contradicts the idea that the mechanism of multitolerance is primarily because of metabolic inactivity [2, 23]. Genes in Toxin-Antitoxin modules were also differentially regulated during persistence, but some were upregulated while others were down-regulated. For example, the TisB toxin was down-regulated during spontaneous persistence, despite the fact that it has been shown that TisB can induce persistence [2, 13]

Several genes related to stress response mechanisms were down-regulated, whereas only two stress-related genes, *zraP* and *pspD*, were up-regulated. This regulation in spontaneous persistence was contrary to previous studies on triggered persistence [2, 8, 23, 28]. Notably, all genes related to the SOS response showed no transcriptomic change in expression or were beyond our resolution power. This particular and striking difference between the current expression profile of spontaneous persister cells and all previous transcriptomic studies performed on persister cells could be caused by the inherent differences between spontaneous and triggered persister cells. Notably, previous studies routinely isolate persisters by using antibiotics e.g. fluoroquinolones that induce the SOS response. The induction of the SOS response by antibiotics during the isolation of persister cells in all previous studies is likely to have caused significant alterations to the expression profiles of the isolated persisters. Our findings suggest that the induction of stress mechanisms, such as the stringent response, acidic shift and oxidative stress, only cause persistence induction if the SOS response has been previously induced, as suggested by the absence of these mechanism in the set of upregulated genes in spontaneous persisters

Gene expression information could significantly aid in the discovery of the genes responsible for the high persistence phenotype of DS1. However, expression patterns can be hard to interpret, as they might be causative, responsive, or independent of persistence. But by correlating the discovered SNPs in the genome of *E. coli* DS1 with transcriptome data, we propose 28 candidate genes that may be involved somehow in persistence (Table 4). For example, the outer membrane porin *ompF*, which is known to mediate the entry of various antibiotics [53, 54] was found to have suffered a deletion which resulted in a predicted frame shift, which might severely affect the functionality of this porin. Another interesting candidate mutation was found in the MltG protein (formerly *yceG*). MltG is an inner membrane enzyme with endolytic murein transglycosylase activity which has been proposed to terminate nascent peptidoglycan synthesis [55] and its deletion has been reported to reduce sensitivity to ampicillin [56]. Additionally, the non-synonymous mutations found in the genes *rho, rpoB, rseB, hflX* and *rsxG* could broadly affect gene expression via alteration of transcription factor activity, translation regulation and mRNA stability, which could also increase noise in gene expression favoring the entrance into alternate phenotypic states.

A notable result of our transcriptomics analysis was the overrepresentation of genes related to fatty acid metabolism and lipid modifications among the biological functions strongly up-regulated during spontaneous persistence. These results not only correlate with some of the mutations found in the DS1 genome, but also support previous findings about differences in the cell membrane of persister cells [24]. The chemical structure of fatty acids determines the level of lipid packing, and influences the biophysical properties of bacterial membranes [57]. We therefore analyzed the fatty acid composition of triggered and spontaneous persister cells and compared the composition with normally growing cells from the wild type strain *E. coli* MG1655 in stationary phase and from the *E. coli* DS1 strain in exponential growth, respectively. The fatty acid composition of persister cells differed significantly from that of the normal cells. Specifically, triggered persisters isolated during stationary phase display a likely increase in the overall membrane rigidity, while spontaneous persisters isolate during exponential growth phase appear to increase their membrane fluidity. The regulation of the physicochemical properties of the cell membrane is a known bacterial strategy to survive challenging environments [57]. Previous studies have also shown a relationship between membrane fluidity and tolerance to certain antibiotic agents [58]. Therefore, alterations in membrane composition and fluidity emerge as a strong candidate for the ultimate mechanism of multitolerance.

## Conclusions

In summary, we combined an isolation protocol for spontaneous persisters [24] with multi-omics methods, to characterize the high persistence strain DS1 hipQ in terms of polymorphisms as well as broad changes in the transcriptome and lipid composition. This revealed many striking changes, excluded previously proposed gene and locus claimed to cause persistence, and identified a set of new candidate genes. Finally, we noted differences in the membrane lipid composition between persister and non-persister cells, thus opening a new avenue of research on the biochemical mechanism of multitolerance.

## Methods

### Bacterial strains and growth conditions

The bacterial strains used in this work and their relevant characteristics are described in Table 4. Bacterial cultures were grown in Luria-Bertani (LB) medium at 37°C and 200 rpm unless otherwise specified. For exponential growth, samples were taken at OD600 = 0.4. The morphological characteristics of strains DS1, KL16 and MG1655 were analyzed at this point and are depicted in Additional file 9.

**Table 4.** *Escherichia coli* strains used in this study.

| Strains | Relevant features | Source or reference |
| --- | --- | --- |
| TH1268 | MG1655 zde264::Tn10 | Korch et al, 2003 |
| TH1269 | MG1655 *hipA7* zde264::Tn10 | Korch et al, 2003 |
| DS1 | *hipQ* | Wolfson et al, 1990 |
| DSY | *hipQ* with *yfp-Cam* cassette | Fridman et al., 2014 |
| KL16 | K-12 $\lambda^-$, *e14-*, *relA1*, *spoT1*, *thiE1* | Low B., 1968 |
| MG1655 | MG1655 6300 |  |
| MG1655 $\Delta ydcI$ | MG1655 6300 $\Delta ydcI$ | This study |

### Escherichia coli DS1 genome sequencing, assembly and annotation

Total DNA was purified from a stationary phase culture of *E. coli* DS1 using the GenElute Bacterial Genome Extraction Kit (Sigma-Aldrich, St. Louis, USA) according to the manufacturer’s protocols. The DS1 genome was sequenced on an Illumina HiSeq 2000 instrument using the 2×90 paired-end technology with a 500 bp insert size at BGI Genomics (Shenzhen, China).

FastQC (Babraham Bioinformatics, Cambridge, United Kingdom) was utilized to visually inspect quality metrics of the raw reads. Reads were clipped, quality trimmed and quality filtered (with a minimum read length of 60 bps and a quality threshold of 20) using Flexbar [59].

Clean reads were then *de novo* assembled using the CLC Assembly Cell (CLC Bio, Aarhus, Denmark). PET scaffolding was performed using the SSAKE-based Scaffolding of Pre-Assembled Contigs after Extension (SSPACE) v2.0 [34]. The PAGIT (Post Assembly Genome Improvement Toolkit) [33] was utilized for reference-guided contig extension using ABACAS [60], PET gap closing was performed using IMAGE [61] and the quality assessment of the assembly was made using iCORN [62]. Finally, Gapfiller was used [34] to close the majority of the remaining gaps. DS1 genome annotation was performed using the Rapid Annotations using Subsystems Technology program [35]. For visualization, a circular representation of the DS1 genome was created using DNAplotter [36].

### Whole genome SNP detection

SNP calling was performed as described by the BROAD GATK Best practices guidelines (v3) using both *E. coli* MG1655 and DH10B genomes as references. Briefly, the sequenced small reads were mapped against each reference genome using the Burrows-Wheeler Aligner (BWA) [63], and the coverage depth was analyzed using the Genome Analysis Toolkit (GATK) [64], obtaining a 94X coverage for each genome. Next, duplicates were marked using Picard [http://picard.sourceforge.net] before performing local re-alignments. For the base quality recalibration step, we built a database of polymorphic sites in *E. coli* using the genomes of strains BL21, S88, 0127:H6 E2348/69, O42, ETEC H10407, DH10B and MG1655, employing progressive Mauve [65] for the multiple sequences alignment. Finally, SNPs in the genome of *E. coli* DS1 were called against each reference genome with GATK [66]

### Analysis of overrepresented GO terms in set genes encoding novel mutations in E. coli DS1

The set of genes identified to have novel mutations, with respect to the three reference strains, was singularly mapped to UniProt accession identifiers using PANTHER [39, 40]. The 95 unique entries were then analyzed for overrepresentation of Cellular component GO terms using DAVID 6.7 [41, 42] with the Fisher’s exact test, with a false discovery rate (FRD) correction to account for multiple testing.

### RNASeq analysis of spontaneous persister cells from exponentially growing E. coli K12 DS1

To obtain enough RNA from the spontaneous persister cells for the RNASeq analysis, 6 replicate flasks, each containing 300 mL of LB media, were individually inoculated with 10 μL of an overnight culture of *E. coli* DS1 and incubated at 37LJC and 200 rpm until the culture reached an OD of 0.4. After reaching the desired OD, the bacterial cultures were pelleted, immediately frozen and stored at -30LJC. This was repeated until a total of 9.3 liters of exponentially grown cultures was processed for each of the two biological replicates.

To extract total RNA from spontaneous persister cells, persisters were first isolated by employing the lysis protocol described in [24], and the RNA from the lysed non-persister cells was completely degraded using RNAse A (Sigma-Aldrich, St. Louis, USA) before proceeding with the total RNA extraction. The complete degradation of the RNA from the non-persister cells was determined with a gel electrophoresis of RNA extractions of the supernatant. Prior to the extraction of the total RNA from persisters, the pellets were washed three times.

Phenol-chloroform RNA extractions were performed in duplicate for both exponentially growing cells and DS1-(*hipQ*)-strain spontaneous persister cells from a pellet equivalent to an initial culture of 150 mL and 4.5 L for each biological replicate, respectively. DNA degradation and total RNA purification were performed with a Qiagen RNeasy kit according to the manufacturer’s protocols (Qiagen, Hilden, Germany). RNASeq on each sample was performed at BGI Genomics on an Illumina HiSeq 2000 instrument using 2×101 paired-end tags and strand-specific chemistry. Raw reads were processed as indicated above.

### Transcriptome assembly, annotation, and analysis

For spontaneous persisters and exponentially growing cells from the DS1 strain, we assembled the complete transcriptome using Trinity [67] for both *de novo* and genome-guided assemblies. Each transcriptome was then used for protein prediction and annotation of genes using Trinotate [67]. The Trinotate pipeline includes a homology search to known sequence data using blastx and blastp [68], the identification of protein domains using HMMER v3.0 [69], a prediction of signal peptides with singalP [70] and tmHMM [71], and several comparisons to curated annotation databases, such as EMBL, Uniprot, KEGG [72], eggNOG [73], and Gene Ontology [74].

TopHat2 was employed to map all PET reads to the reference genome [75]. Expression levels were presented as Fragments per Kilobase of exon per Million reads (FPKMs), and differential gene expression analyses were performed using both CuffDiff2 [76, 77] and NOISeq [78] using the cutoffs of a p-value of 0.05 and a q of 0.9, respectively.

### Analysis of overrepresented GO terms in differentially expressed genes during spontaneous persistence

To analyze the biological processes that are significantly regulated during spontaneous persistence, we performed a Blast2GO [46] analysis over the complete *E. coli* deduced proteome. We then tested for significant overrepresentation of GO terms in the groups of differentially expressed, overexpressed and underexpressed genes derived from CuffDiff [76]and NOISeq [78] analysis performed between *E. coli* DS1 exponentially growing cells and spontaneous persisters. The overrepresentation analysis was performed using DAVID 6.7 [41, 42] was then used to test for overrepresentation with the Fisher’s exact test, with a false discovery rate (FRD) correction to account for multiple testing (Benjamini-Hochberg test) and Bonferroni’s score both with a threshold of <0.005.

### RT-qPCR validation of detected differentially expressed genes

From the group of differentially expressed genes, the expression profiles of 14 genes were chosen to be validated with RT-qPCR. The selection of these genes accounted for their biological function and the existence of previous reports of their relevance to triggered persistence [2, 8, 28]. The housekeeping genes *dxs* and *opgH* were used to normalize the data. For this analysis, a sample of the identical RNA that was sequenced and the RNA from a biological replica for each condition were converted to cDNA using random hexamers prior to the qPCR. The cDNA synthesis and qPCR were performed with the DyNAmo SYBR Green 2-Step qRT-PCR Kit (Thermo Scientific, Waltham, USA).

The validation of the differentially expressed genes through qPCR was performed using the relative quantification method with a standard curve on a 7500 Fast qPCR Instrument (Applied Biosystems, Life Technologies, California, USA). The statistical analysis of the obtained data was done with REST [79] using the Pair Wise Fixed Reallocation Randomization Test method [79].

### Determination of total fatty acids

Lipids were extracted from *E. coli* strains MG1655 and DS1 during exponential growth and stationary phase. Triggered and spontaneous persister cells were isolated from a stationary phase culture of *E. coli* TH1269 and an exponential phase culture of *E. coli* DS1, respectively, using a published protocol [24].

For each of the above-mentioned samples, lipids were extracted by pelleting 60 mL of each culture (or its equivalent in cell population for the triggered and spontaneous persister cells after the isolation protocol); the pellets were then frozen at -20^0^ C for 2 hours. Frozen pellets were then lyophilized overnight to remove any residual water. Afterwards, samples were dispersed in a chloroform/methanol/water (3:1:1 v/v) mixture and then vortexed every 15 minutes for 4 hours.

After two days three separate phases are visually identified, where the top phase is an aqueous phase, the middle phase is a protein-rich phase, and the lower phase is the organic phase enriched in total lipids. The lower organic phase was drawn off by aspiration and collected into clean glass tube. All glass is cleaned using a piranha (sulfuric acid and hydrogen peroxide) protocol to remove all organic residues. Chloroform was then evaporated with a steady stream of gaseous N_2_ to form a thin film at the bottom of the test tube, and the lipids were then stored at -20C. Extracted lipids were analyzed by gas chromatography.

Preparation of methyl esters of fatty acids (FAMEs) for analysis by gas chromatography/flame ionization detection (GC/FID) was performed as already described [80]. For acidic hydrolysis, 1 ml methanol/toluene (2:1, v/v) containing 2.75 % (v/v) H_2_SO_4_ (95-97 %) and 2 % (v/v) dimethoxypropan was added to the dry sample material. For later quantification of the fatty acids, 20 µg triheptadecanoate (Tri-17:0) were added and the sample was incubated for 1 h at 80 °C. To extract the resulting FAMEs, 200 µl of saturated aqueous NaCl solution and 2 ml of hexane were added. The hexane phase was dried under streaming nitrogen and re-dissolved with equal volumes of water and hexane. The hexane phase was filtrated with cotton wool soaked with NaSO_4_ and dried under streaming nitrogen. Finally, the sample was re-dissolved in 10 µl acetonitrile for GC analysis performed with a Agilent (Waldbronn, Germany) 6890 gas chromatograph fitted with a capillary DB-23 column (30 m x 0.25 mm; 0.25 µm coating thickness; J&W Scientific, Agilent). Helium was used as carrier gas at a flow rate of 1 ml/min. The temperature gradient was 150°C for 1 min, 150 – 200°C at 8 K/min, 200–250°C at 25 K/min and 250°C for 6 min. For verification of the peak identities, an aliquot of the sample was analysed by GC/ mass spectrometric detection (GC/MS) using an Agilent 5973 Network mass selective detector connected to the gas chromatograph as described above. The injection temperature was 220°C. The temperature gradient as well as the carrier gas was carried out as described for the GC analysis. Electron energy of 70 eV, an ion source temperature of 230 °C, and a temperature of 350°C for the transfer line were used. Mass detection was performed in scan mode in an *m*/*z* range of 50 to 650. Lipid extraction and analysis were performed for each of the above-described samples with three biological replicates.

## Declarations

### Ethics approval and consent to participate

Not applicable

### Consent for publication

Not applicable

### Availability of data and materials

All the genomic sequence data generated for the *E. coli* DS1 (hipQ) has been submitted to the SRA as the entry accessioned SRX589587. The assembled and the annotated genome can be accessed in GenBank (CP022466). All the data from the transcriptomic analyzes of type I and type II persisters has also been submitted to SRA with accession numbers PRJNA252678 and PRJNA253803.

### Authors’ contributions

S.J.C., J.M.P. and S.R. designed research: S.J.C., L.S., M.I.P., C.H., L.M.C. and D.M.R performed the research; I.F., C.L., D.M.R., J.P., J.M.P. and S.R. contributed new reagents/analysis tools; S.J.C., L.S., M.I.P., C.L., D.M.R., J.P., J.M.P. and S.R. analyzed data; S.J.C., L.S., M.I.P., C.L., D.M.R., J.P., J.M.P. and S.R. wrote the paper.

## Acknowledgements

We thank the laboratory of D. C. Hooper for retrieving the E. coli DS1 (hipQ) strain. We also thank Dr. Thomas M. Hill for sending us strains TH1268 and TH1269. We further thank the Laboratory of Nathalie Balaban for sending us the strains DSY and KLY.

