## Additional file 2 for "An integrative multi-omics approach points to membrane composition as a key factor in *E. coli* persistence"

Title: Prediction of *E. coli* DS1 HipA protein aligned to the crystal structure of HipA of MG1655

Legend: AlphaFold2 prediction made using ColabFold v1.5.2 [1] of the HipA protein carrying the novel A242V mutation identified in the genome of *E. coli* DS1. The resultant structure (silver) was aligned to the crystal structure of HipA from MG1655 [2] (blue) using the Pairwise Structure Alignment tool from RCSB PDB [3]. The model for DS1's HipA is depicted in silver, and MG1655 HipA is depicted in blue. A zoomed in view into the region of residue 242 is shown as an inset.

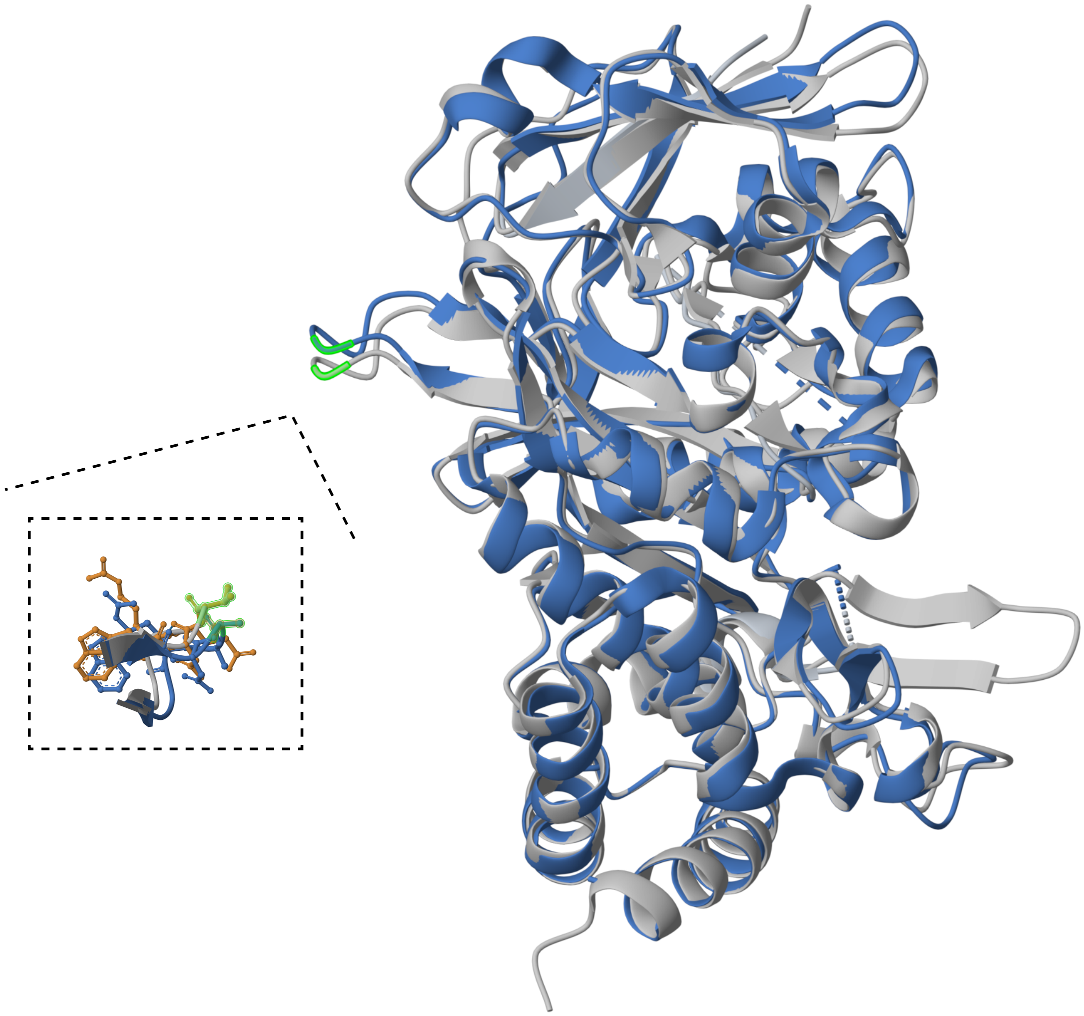
