## Additional file 3 for "An integrative multi-omics approach points to membrane composition as a key factor in *E. coli* persistence"

Title: Schematic of persister isolation protocol using antibiotics or the lysis protocol

Legend: Traditional antibiotic-based protocols typically involve a short pre-growth phase (e.g., 2 hours for 100x diluted stationary cultures) in order to efficiently kill stationary phase cells, which is followed by an extended antibiotic treatment (3–5 hours). This prolonged treatment may induce stress in spontaneous persisters, potentially altering their cellular components. Additionally, it may trigger normal cells to transition into triggered persisters. In contrast, our lysis-based protocol incorporates an extended pre-growth phase followed by rapid lysis-based isolation, mitigating these confounding variables.

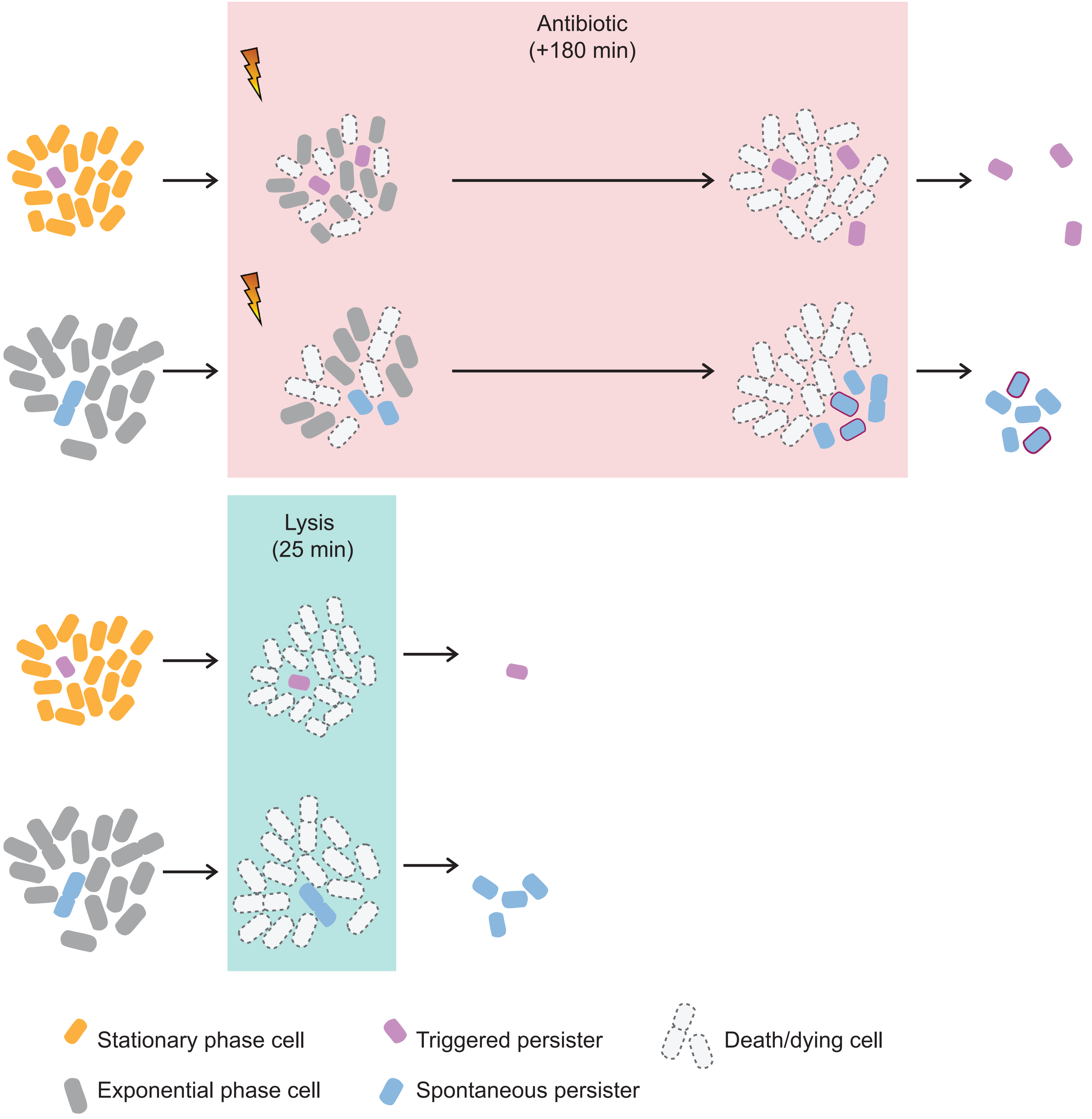
