## Additional file 6 for "An integrative multi-omics approach points to membrane composition as a key factor in *E. coli* persistence"

Title: RT-qPCR validation results
Legend: Normalized and non-normalized calculations of the relative expression of the 12 differentially expressed genes chosen for validation.

### Relative Expression Results

| Parameter | Value |
|---|---|
| Iterations | 6000 |

| Gene | Type | Reaction Efficiency | Expression | Std. Error | 95% C.I. | P(H1) Result |
|---|---|---|---|---|---|---|
| opgH | REF | 1.2923 | 0.764 | | | |
| dxs | REF | 0.7966 | 1.309 | | | |
| fadB | TRG | 0.8406 | 5.047 | 2.881 - 10.588 | 1.873 - 12.745 | 0.035 UP |
| tisB | TRG | 0.7683 | 0.191 | 0.104 - 0.512 | 0.059 - 0.631 | 0.035 DOWN |
| cadA | TRG | 1.1603 | 0.058 | 0.033 - 0.110 | 0.022 - 0.140 | 0.035 DOWN |
| relB | TRG | 0.9881 | 0.172 | 0.118 - 0.319 | 0.067 - 0.347 | 0.035 DOWN |
| dinJ | TRG | 0.6368 | 0.401 | 0.270 - 0.661 | 0.192 - 0.760 | 0.019 DOWN |
| soxS | TRG | 0.6951 | 4.951 | 3.271 - 8.067 | 2.935 - 9.879 | 0.035 UP |
| hde | TRG | 0.6977 | 0.109 | 0.066 - 0.161 | 0.056 - 0.192 | 0.024 DOWN |
| acs | TRG | 0.6848 | 2.676 | 1.873 - 4.125 | 1.455 - 4.937 | 0.035 UP |
| iraP | TRG | 0.6507 | 0.263 | 0.132 - 0.539 | 0.081 - 0.629 | 0.035 DOWN |
| ompX | TRG | 0.9019 | 0.125 | 0.067 - 0.182 | 0.060 - 0.243 | 0.035 DOWN |
| pspB | TRG | 1.5741 | 22.295 | 11.716 - 42.509 | 8.006 - 50.970 | 0.025 UP |
| groS | TRG | 1.5722 | 0.155 | 0.125 - 0.195 | 0.108 - 0.230 | 0.035 DOWN |
| yodD | TRG | 1.4021 | 0.222 | 0.148 - 0.450 | 0.075 - 0.480 | 0.035 DOWN |
| tnaA | TRG | 1.5454 | 8.520 | 3.767 - 15.132 | 2.890 - 32.011 | 0.013 UP |

**Interpretation**

fadB is UP-regulated in sample group (in comparison to control group) by a mean factor of 5.047 (S.E. range is 2.881 - 1
fadB sample group is different to control group. P(H1)=0.035

tisB is DOWN-regulated in sample group (in comparison to control group) by a mean factor of 0.191 (S.E. range is 0.104
tisB sample group is different to control group. P(H1)=0.035

cadA is DOWN-regulated in sample group (in comparison to control group) by a mean factor of 0.058 (S.E. range is 0.03
cadA sample group is different to control group. P(H1)=0.035

relB is DOWN-regulated in sample group (in comparison to control group) by a mean factor of 0.172 (S.E. range is 0.118
relB sample group is different to control group. P(H1)=0.035

dinJ is DOWN-regulated in sample group (in comparison to control group) by a mean factor of 0.401 (S.E. range is 0.270
dinJ sample group is different to control group. P(H1)=0.019

soxS is UP-regulated in sample group (in comparison to control group) by a mean factor of 4.951 (S.E. range is 3.271 - [illegible]
soxS sample group is different to control group. P(H1)=0.035

hde is DOWN-regulated in sample group (in comparison to control group) by a mean factor of 0.109 (S.E. range is 0.066
hde sample group is different to control group. P(H1)=0.024

acs is UP-regulated in sample group (in comparison to control group) by a mean factor of 2.676 (S.E. range is 1.873 - 4.[illegible]
acs sample group is different to control group. P(H1)=0.035

iraP is DOWN-regulated in sample group (in comparison to control group) by a mean factor of 0.263 (S.E. range is 0.132
iraP sample group is different to control group. P(H1)=0.035

ompX is DOWN-regulated in sample group (in comparison to control group) by a mean factor of 0.125 (S.E. range is 0.0[illegible]
ompX sample group is different to control group. P(H1)=0.035

pspB is UP-regulated in sample group (in comparison to control group) by a mean factor of 22.295 (S.E. range is 11.716
pspB sample group is different to control group. P(H1)=0.025

groS is DOWN-regulated in sample group (in comparison to control group) by a mean factor of 0.155 (S.E. range is 0.12[illegible]
groS sample group is different to control group. P(H1)=0.035

yodD is DOWN-regulated in sample group (in comparison to control group) by a mean factor of 0.222 (S.E. range is 0.14
yodD sample group is different to control group. P(H1)=0.035

tnaA is UP-regulated in sample group (in comparison to control group) by a mean factor of 8.520 (S.E. range is 3.767 - 1 tnaA sample group is different to control group. P(H1)=0.013

#### Non-Normalised Results

| Gene | Type | Reaction Efficiency | Expression | Std. Error | 95% C.I. | P(H1) | Result |
|---|---|---|---|---|---|---|---|
| opgH | REF | 1.2923 | 0.813 | 0.677 - 0.978 | 0.607 - 1.112 | 0.263 | |
| dxs | REF | 0.7966 | 1.392 | 0.636 - 3.618 | 0.462 - 5.107 | 0.465 | |
| fadB | TRG | 0.8406 | 5.369 | 4.631 - 6.579 | 4.343 - 7.045 | 0.000 | UP |
| tisB | TRG | 0.7683 | 0.203 | 0.118 - 0.336 | 0.116 - 0.346 | 0.048 | DOWN |
| cadA | TRG | 1.1603 | 0.062 | 0.053 - 0.071 | 0.051 - 0.081 | 0.000 | DOWN |
| relB | TRG | 0.9881 | 0.183 | 0.144 - 0.231 | 0.137 - 0.245 | 0.000 | DOWN |
| dinJ | TRG | 0.6368 | 0.427 | 0.382 - 0.462 | 0.370 - 0.472 | 0.000 | DOWN |
| soxS | TRG | 0.6951 | 5.267 | 4.210 - 6.829 | 3.881 - 7.648 | 0.048 | UP |
| hde | TRG | 0.6977 | 0.116 | 0.102 - 0.143 | 0.099 - 0.147 | 0.000 | DOWN |
| acs | TRG | 0.6848 | 2.847 | 2.337 - 3.452 | 2.282 - 3.504 | 0.035 | UP |
| iraP | TRG | 0.6507 | 0.279 | 0.214 - 0.362 | 0.182 - 0.373 | 0.030 | DOWN |
| ompX | TRG | 0.9019 | 0.133 | 0.107 - 0.152 | 0.103 - 0.185 | 0.010 | DOWN |
| pspB | TRG | 1.5741 | 23.717 | 15.682 - 30.956 | 14.412 - 49.322 | 0.000 | UP |
| groS | TRG | 1.5722 | 0.164 | 0.100 - 0.277 | 0.076 - 0.378 | 0.000 | DOWN |
| yodD | TRG | 1.4021 | 0.236 | 0.182 - 0.348 | 0.149 - 0.388 | 0.000 | DOWN |
| tnaA | TRG | 1.5454 | 9.063 | 5.878 - 18.211 | 3.065 - 20.524 | 0.000 | UP |
