## Additional file 9 for "An integrative multi-omics approach points to membrane composition as a key factor in *E. coli* persistence"

Title: Morphological characteristics of exponentially growing *E. coli* strains DS1, KL16 and MG1655 at OD600 =0.4

Legend: Cultures were grown in 50 mL of LB at 37C in 250 mL flasks until they reached an OD600 of 0.4. Samples were spotted in 1.5% LB-agarose pads and imaged using a 100x Ph3 objective in a Nikon Ti2 inverted microscope at 37C. Supersegger-Omnipose [1-3] was used to segment the cells and an inhouse Matlab code was used to analyze the extracted features.

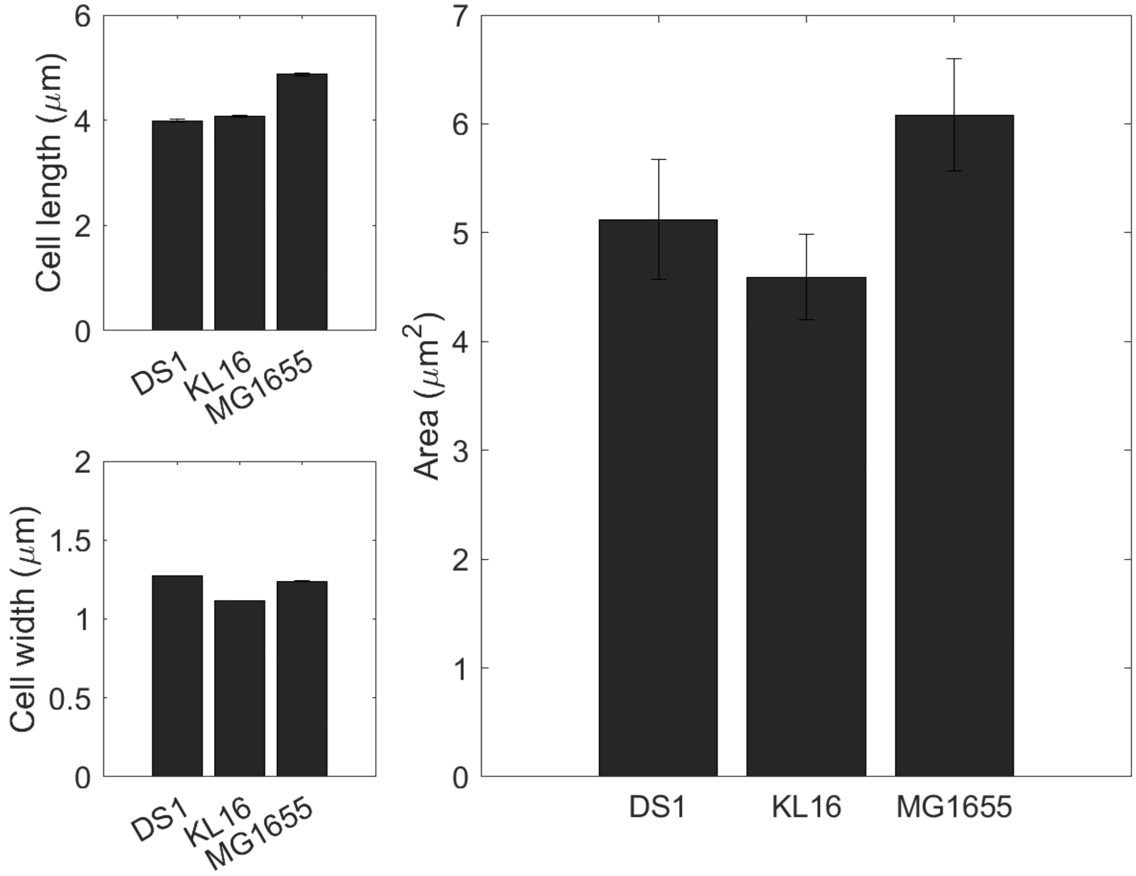
